# State dependent shifts in large scale functional topographies

**DOI:** 10.1101/2025.10.16.682912

**Authors:** Yezhou Wang, Jordan DeKraker, Raul Rodriguez Cruces, Donna Gift Cabalo, Jessica Royer, Alexander Ngo, Meaghan Smith, Brontë McKeown, Youngeun Hwang, Ilana Leppert, Tamara Vanderwal, R. Nathan Spreng, Sofie L. Valk, Jonathan Smallwood, Alan C Evans, Boris Bernhardt

## Abstract

Although functional networks can be consistently identified across cognitive states, they also undergo dynamic reconfigurations across different contexts. For example, naturalistic movie watching paradigms amplify activity in sensory systems compared to resting conditions. However, it remains unclear how these different states affect large-scale brain organization. The current study leveraged high-resolution *in vivo* 7T fMRI data from the Human Connectome Project (HCP) and the Precision NeuroImaging (PNI) datasets to examine large scale functional connectivity changes between resting and movie-watching conditions. To understand these changes within topographic and geometric principles of brain organization, connectivity shifts were stratified relative to macroscale cortical hierarchy and geodesic distance. Our results revealed that primary sensory areas showed increased local connectivity and reduced long-range interactions during movie watching relative to resting conditions, whereas the default mode network (DMN) exhibited an opposing pattern characterized by reduced within-network long-range connectivity and enhanced connectivity with distant regions outside the DMN. Together, these findings demonstrate that different cognitive states involve geometry-and hierarchy-informed reorganization of large-scale functional networks.

## Introduction

In recent decades, resting-state functional MRI (rs-fMRI) has become a key method to study large-scale functional organization of the human brain (Biswal et al., 1995; Fox et al., 2005; Raichle et al., 2001; Smith et al., 2009). By measuring spontaneous fluctuations in brain activity during wakeful rest, and their inter-regional statistical dependencies, rs-fMRI has revealed consistent and reproducible large-scale networks, such as the default mode, frontoparietal, salience, and sensory networks (Glasser et al., 2016; Power et al., 2011; Yeo et al., 2011). These networks form part of a macroscale topography that reflects the brain’s functional architecture and supports the integration of information across distinct cognitive domains. Beyond identifying networks, rs-fMRI studies have also revealed gradual spatial patterns that organize the cortex along hierarchical axes, from unimodal sensory regions to heteromodal association areas (Bernhardt et al., 2022; Margulies, Ghosh, et al., 2016; Mesulam, 1998). This organization is thought to support increasing levels of abstraction and multimodal integration, enabling adaptive cognition in complex environments. Importantly, functional connectivity patterns have been shown to correlate with anatomical features such as geodesic distance, which reflects the shortest path along the cortical surface and approximates spatial wiring cost (Alexander-Bloch et al., 2013; Goulas et al., 2017). Meanwhile, hierarchical models posit that cortical processing unfolds along continuous gradients from low-level sensory to high-level association regions, with functional heterogeneity increasing along this axis (Bernhardt et al., 2022; Demirtaş et al., 2019; Huntenburg et al., 2018; Paquola et al., 2019; Smallwood et al., 2021; Sydnor et al., 2021). Together, these findings point to a close link between intrinsic functional organization, cortical hierarchy, and anatomical geometry.

Rs-fMRI has been instrumental in mapping the brain’s intrinsic architecture, and there is furthermore work showing a close relationship to brain activation and connectivity patterns during laboratory tasks (Cole et al., 2016; Ngo et al., 2022; Smith et al., 2009; Tavor et al., 2016). On the other hand, we still lack an in-depth understanding on whether and how functional organization reconfigures during dynamic, ecologically valid conditions. Such settings can be approximated using naturalistic paradigms, such as movie-watching. These paradigms have gained popularity to probe brain function under structured, temporally extended stimulation (Hasson et al., 2004; Sonkusare et al., 2019). Naturalistic stimuli simultaneously engage multiple sensory and cognitive systems, eliciting reliable, widespread brain responses across individuals. Prior work has shown that movie-watching fMRI leads to higher test–retest reliability, stronger inter-subject synchronization, and improved prediction of individual traits relative to rest (Finn et al., 2015; Vanderwal et al., 2017). These paradigms have been shown to amplify individual differences in functional architecture and better capture behavioral and cognitive profiles (Finn & Bandettini, 2021; Vanderwal et al., 2019). Moreover, movie-watching tasks broadly engage the cortical hierarchy, from early sensory cortices to transmodal areas involved in memory, language, and social cognition (Nastase et al., 2020). Although prior studies have examined how functional connectivity patterns shift between resting and naturalistic states (Samara et al., 2023), relatively little is known about how these changes relate to cortical processing hierarchies (Bernhardt et al., 2025; Wei et al., 2024) and geometric constraints of brain function (Pang et al., 2023).

The goal of this study is to investigate how functional connectivity reconfigures during naturalistic stimulation and how these changes are constrained by cortical hierarchy and anatomical distance. To this end, we leveraged high-resolution *in vivo* multistate 7T fMRI data from the HCP (Van Essen et al., 2013) and the PNI (Cabalo et al., 2025) datasets to investigate connectivity alterations between resting-state and movie-watching conditions, with a focus on their relationships to cortical processing hierarchies and geometry. The inclusion of two independent datasets allowed us to assess the generalizability and robustness of our findings across different movie stimuli, acquisition protocols, and participant cohorts, while benefiting from enhanced resolution, signal-to-noise ratio, and anatomical precision of 7T MRI. We focused on three representative regions of interest (ROIs) that span the unimodal–transmodal hierarchy and quantified the movie–rest difference (MRD) in functional connectivity patterns for each region. We then examined how MRD values varied across large-scale functional networks and established hierarchical levels, defined using foundational accounts of functional hierarchies in the primate cortex (1998). Importantly, we investigated whether geodesic distance between ROIs predicted the magnitude and direction of functional connectivity changes between rest and movie-watching conditions, and whether these relationships differed across unimodal and transmodal areas. By combining naturalistic fMRI with cortical hierarchy models and geometric constraints, our study aims to advance understanding of the principles that govern functional reorganization across brain states.

## Methods

### Participants

*HCP-*7T (Van Essen et al., 2013). For our main analysis, we studied rs-fMRI and movie-watching fMRI data from 93 healthy adults (mean age 29.4±3.3 years, 56 females) from the HCP 7T dataset (Glasser et al., 2013). Each participant underwent four sessions on separate days. All data was collected as part of the Human Connectome Project under protocols approved by the Washington University Institutional Review Board, and informed consent was obtained from all participants. *MICA-PNI* (Cabalo et al., 2025). To validate our findings, we repeated all analyses in an independent 7T dataset with 30 unrelated healthy adults (age: 29.20±5.20 years, 15 females). Each participant underwent three sessions on separate days. Data was collected between March 2022 and June 2024. The first data release including data from 10 individuals is openly available at the OSF platform (https://osf.io/mhq3f/). The studies were approved by the Ethics Committees of McGill University and the Montreal Neurological Institute and Hospital, respectively, and written and informed consent was obtained from all participants.

### MRI acquisition

*HCP-7T.* The HCP-7T dataset (Van Essen et al., 2013) includes high-resolution MRI data collected on a 7T Siemens Magnetom scanner with a Nova32 head coil at the Center for Magnetic Resonance Research at the University of Minnesota. T1-weighted (T1w) structural images were acquired using a 3D MPRAGE sequence with parameters optimized for ultra-high field imaging. The acquisition parameters included 0.7 mm isotropic voxels, matrix=320×320, 224 sagittal slices, repetition time (TR)=2400 ms, echo time (TE)=2.14 ms, inversion times (TI)=1000 ms, flip angles=8°, and bandwidth=210 Hz/px. Resting-state and movie-watching fMRI data were acquired using a gradient-echo planar imaging (EPI) sequence with the following parameters: TR=1000 ms, TE=22.2 ms, flip angle=45°, field of view (FOV)=208×208 mm², slice thickness = 1.6 mm; 85 slices, multiband factor=5, echo spacing=0.64 ms, bandwidth=1924 Hz/px; 1.6 mm isotropic voxels. Four movie-watching runs varied from 15 minutes 1 second (901 TRs) to 15 minutes 21 seconds (921 TRs) were completed. Each movie run consisted of 4 to 5 distinct clips. Two categories of movies were presented: short independent films released under Creative Commons licenses (Movie 1 and Movie 3), and clips from Hollywood feature films (Movie 2 and Movie 4). Clip durations ranged from 1 minute 4 seconds (64 TRs) to 4 minutes 19 seconds (259 TRs), with a 20-second rest period between clips. All four movie runs concluded with the same 1-minute 24-second (84 TRs) Vimeo video, and each run began and ended with a 20-second rest period. A full description of the imaging protocol and acquisition parameters is available in the HCP 7T data release documentation (https://db.humanconnectome.org/). Four Rest runs were completed, each lasting 15 minutes. During this 15-min rs-fMRI scan, participants were instructed to fixate on a bright crosshair on a dark background and remain still.

*MICA-PNI.* High-resolution T1-weighted structural images were acquired using a 3D MP2RAGE sequence, which produces two images at different inversion times. These were combined to generate quantitative T1 relaxation maps and a bias-corrected synthetic T1w image, offering enhanced myelin sensitivity and reduced B1+ bias (Haast et al., 2016; Marques et al., 2010). Parameters were: 0.5 mm isotropic resolution, matrix size 320 × 320, 320 sagittal slices, TR = 5170 ms, TE = 2.44 ms, flip angle = 4°, TI1/TI2 = 1000/3200 ms, inversion delay = 900 ms, echo spacing = 7.8 ms, bandwidth = 210 Hz/px, partial Fourier = 6/8, and parallel imaging (iPAT) factor = 3. Scans were visually checked for motion artifacts and repeated when necessary to ensure data quality.

Multi-echo fMRI data were acquired using a 2D gradient-echo EPI sequence designed to improve sensitivity and denoising through echo combination. Parameters were: voxel size = 1.9 mm isotropic, TR = 1690 ms, TE1 = 10.8 ms, TE2 = 27.3 ms, TE3 = 43.8 ms, flip angle = 67°, FOV = 224 × 224 mm², slice thickness = 1.9 mm, multiband acceleration factor = 3, and echo spacing = 0.53 ms. During the rs-fMRI scan (∼6 min), participants maintained visual fixation on a central cross and were asked to stay alert while letting their minds wander. Additionally, two spin-echo EPI scans with reversed phase encoding (AP and PA directions) were acquired for geometric distortion correction (TR = 3000 ms, TE = 18.4 ms, flip angle = 90°). Four movie-watching runs were completed under naturalistic viewing conditions to assess brain responses to dynamic audiovisual input (Vanderwal et al., 2017). Two of the runs featured affective movies (Bathroom; Caddy), while the other two consisted of documentary-style movies (Harsh; Pines). Full acquisition protocols are available in the public dataset release (https://osf.io/mhq3f/).

### Multimodal MRI processing

*HCP-7T.* Structural and functional MRI data from the HCP-7T dataset were first processed using the HCP minimal preprocessing pipelines (Glasser et al., 2013), which include gradient unwarping, motion correction, field map-based EPI distortion correction, bias field correction, and surface-based registration to the fsLR-32k space. T1w images underwent surface reconstruction and cortical segmentation via FreeSurfer v6.0 (Fischl, 2012), producing native-space cortical surfaces aligned to individual anatomy and resampled to standard surface space for group-level analysis. All surfaces were visually inspected and manually edited to correct segmentation inaccuracies when necessary.

For rs-fMRI data, we extended the HCP’s FIX-denoised outputs (Glasser et al., 2013; Salimi-Khorshidi et al., 2014) with additional preprocessing steps to ensure compatibility with our analysis pipeline. The initial five volumes of each scan were discarded to allow for signal stabilization. Motion-related artifacts were addressed using spike regression, modeling outlier volumes as nuisance regressors (Lemieux et al., 2007; Satterthwaite et al., 2013). Functional volumes were co-registered to each subject’s cortical surface using boundary-based registration (Greve & Fischl, 2009), as implemented in the HCP pipeline.

*MICA-PNI.* MRI data preprocessing was performed using our multimodal MRI processing pipeline, Micapipe v0.2.3 (Cruces et al., 2022). Structural images were de-obliqued and reoriented, with background denoising (Marques et al., 2010), bias field correction, and intensity normalization (Tustison et al., 2010) applied.. To enhance image contrast, non-local means filtering was applied (Coupe et al., 2008; Tustison et al., 2010), followed by cortical surface reconstruction using FastSurfer v2.2.0 (Henschel et al., 2020). All surface outputs were visually inspected, and additional quality control metrics and manual corrections were used to ensure their accuracy.

Multi-echo resting-and movie-state fMRI data were processed using the same workflow. Each echo was first reoriented and motion corrected using AFNI 23.01.09 (Cox, 1996), after which distortion correction was applied using the reverse phase-encoding scans with FSL TOPUP 6.0.2 (Jenkinson et al., 2012). The scans underwent preprocessing with TEDANA v0.0.12 (DuPre et al., 2021) (https://tedana.readthedocs.io/), integrated within the Micapipe framework (https://micapipe.readthedocs.io/). TEDANA extracted time series from all echoes, optimally combined them, and performed principal component analysis and independent component analysis on the multi-echo BOLD data. Components showing TE dependence were retained as BOLD signal, while non-BOLD components were discarded. Data were then high-pass filtered. Native surfaces were registered from structural to fMRI space via non-linear registration, and volumetric time series were sampled onto these surfaces. Native surface time series were resampled to standard surface templates (fsLR-32k) and the Glasser parcellation (Glasser et al., 2016), followed by motion spike and global signal correction, using linear regression applied to the surface time series.

### Generating functional connectivity for Rest and Movie

*Construction of functional connectivity matrices.* To examine functional connectivity during rest and movie-watching, we first resampled each participant’s fMRI time series from their native surface space to the standard fsLR-32k surface. Parcellation was performed using the Glasser multimodal atlas (Glasser et al., 2016), which divides the cortex into 360 regions based on anatomical and functional features. For each participant, a connectivity matrix was constructed by computing Pearson correlation coefficients between all pairs of regional time series. Correlation values underwent a Fisher’s r-to-z transformation. This process was conducted separately for each of the four movie-fMRI sessions, and resulting matrices were averaged to produce a single movie functional connectivity matrix per subject. The same procedure was applied to the four rs-fMRI runs to generate a corresponding resting-state connectivity matrix.

#### Definition of ROIs within functional networks

To explore hierarchical differences in connectivity reconfiguration, we selected ROIs across twelve canonical functional networks (Ji et al., 2019), ensuring coverage of the cortical hierarchy from primary sensory to transmodal areas. We focused on three networks representing distinct levels of the hierarchy: the visual network (VN), dorsal attention network (dAN), and default mode network (DMN). For each network, we identified the most representative ROI by computing the cosine distance between each parcel’s functional connectivity profile and all other parcels within the same network. The parcel with the lowest mean cosine distance was selected as the ROI, reflecting maximal centrality and representativeness (**Supplementary Figure 1A**). As the result, three ROIs were selected: V1 (VN), MIP (dAN), and 31pv (DMN) (**Figure 1B**).

**Figure 1.**
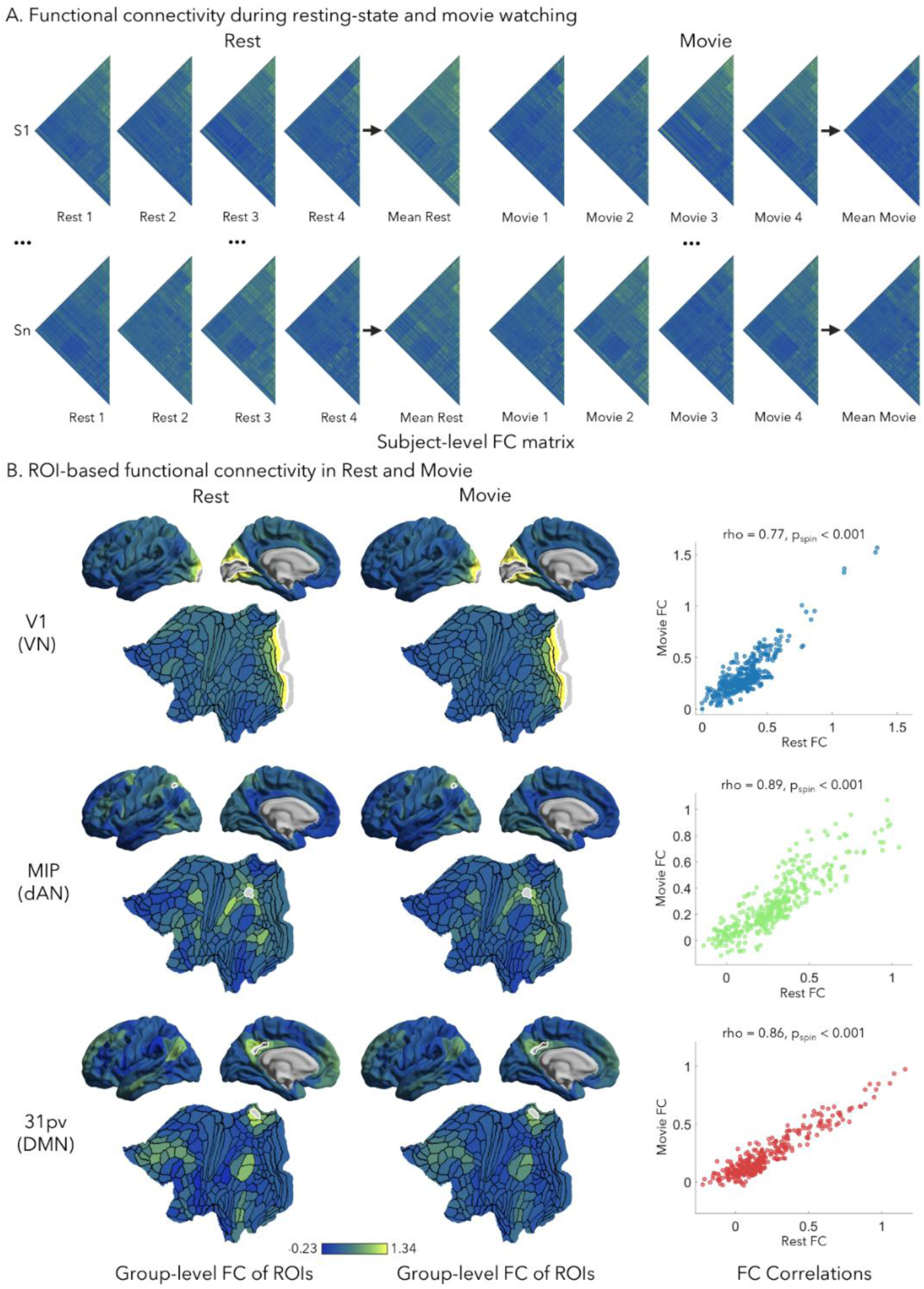
Functional connectivity during resting-state and movie-watching conditions. **(A)** Each participant completed four resting-state and four movie-watching fMRI runs. Time series from each condition were mapped to a multi-modal parcellation (Glasser et al., 2016), and functional connectivity (FC) matrices were computed using Pearson correlation between regional time series. The four FC matrices from each condition were averaged to generate group-level mean Rest FC and mean Movie FC matrices. **(B)** ROI-based FC patterns in Rest and Movie conditions. The left panel illustrates the FC profiles of selected ROIs across the two states, revealing distinct connectivity patterns across functional hierarchies. ROIs outlined in white were chosen as representative nodes within their networks: V1 in the visual network (VN), MIP in the dorsal attention network (dAN), and 31pv in the default mode network (DMN). These ROIs were identified based on their minimal average cosine distance to other regions within each network. The right panel shows the correlation between Rest FC and Movie-watching FC.

**Figure 1.**
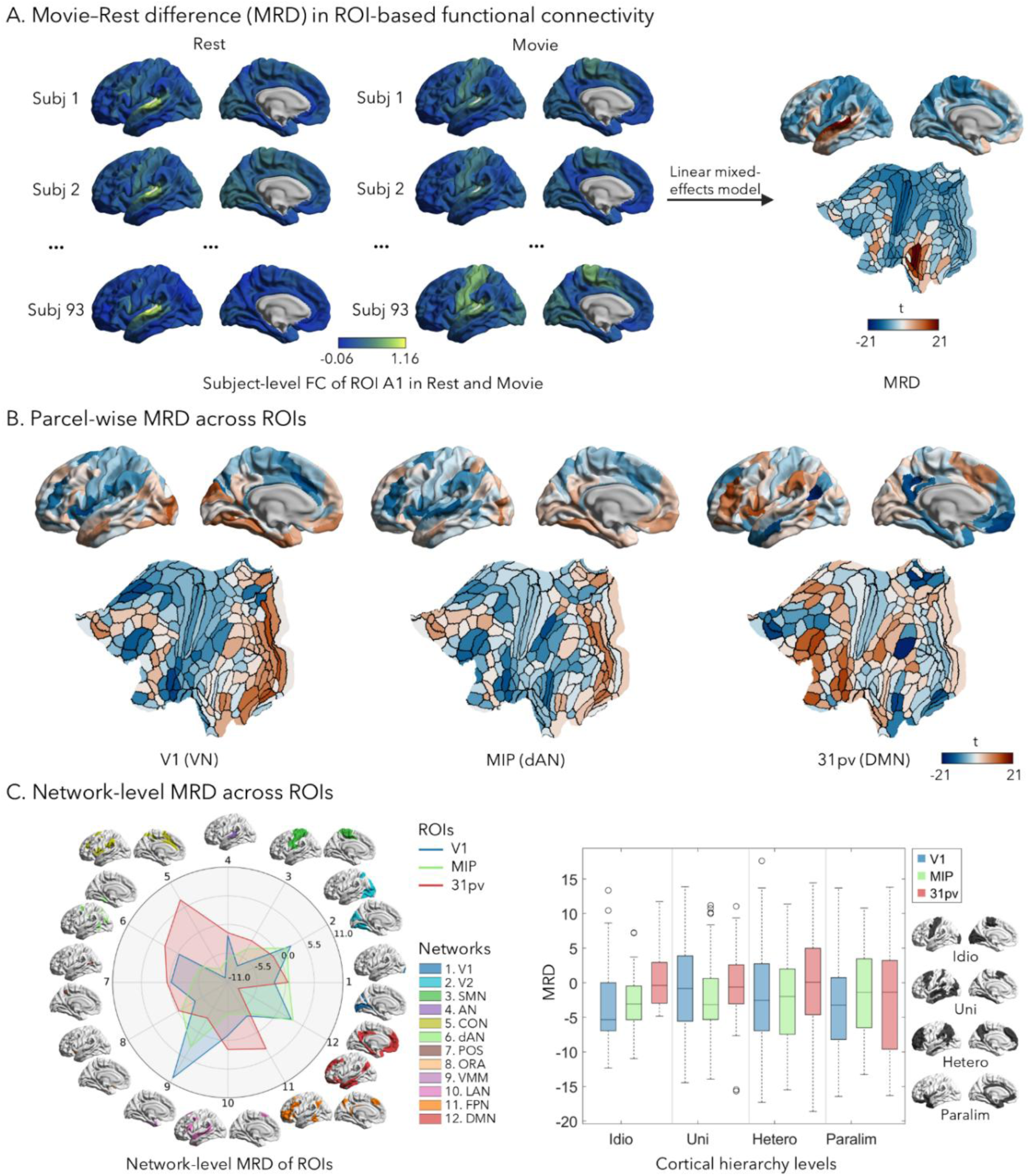
Movie–Rest differences in functional connectivity across spatial scales. **(A)** Movie–Rest Difference (MRD) in ROI-based functional connectivity. For each ROI, MRD was estimated by fitting a linear mixed-effects model to the functional connectivity values across all subjects, with scanning condition (Rest vs. Movie) as a fixed effect and subject as a random intercept. A1 is shown as an example, illustrating how the model captures condition-dependent FC changes relative to all other parcels. **(B)** Parcel-wise MRD across selected ROIs. Parcel-wise MRD maps are displayed for three representative regions: V1 (visual network, VN), MIP (dorsal attention network, dAN), and 31pv (default mode network, DMN). Warmer colors indicate parcels with stronger functional connectivity to the ROI during movie-watching compared to rest. **(C)** Network-level MRD across ROIs. MRD values were aggregated by target network for each ROI to assess distributed reconfiguration patterns across twelve large-scale functional networks and four cortical hierarchy levels (idiotypic, unimodal, heteromodal, paralimbic). V1 showed strong MRD in ventral multimodal areas and low MRD in somatomotor and cingulo-opercular networks. In contrast, 31pv displayed increased MRD in externally oriented networks and reduced MRD within its own DMN, consistent with state-dependent reorganization across cortical hierarchy.

### Estimating movie-rest difference (MRD) in functional connectivity

To quantify how functional connectivity changes between movie-watching and resting-state conditions, we computed the MRD for each ROI. Specifically, a linear mixed-effects model was applied to estimate condition-related differences in functional connectivity across participants (**Figure 2A**). In this model, brain state (rest *vs*. movie) was entered as a fixed effect, while subject identity was treated as a random effect to account for between-subject variability. This approach was used to assess MRD for three representative ROIs, V1, MIP, and 31pv, and the remaining ROIs selected from all twelve functional networks.

**Figure 2.**
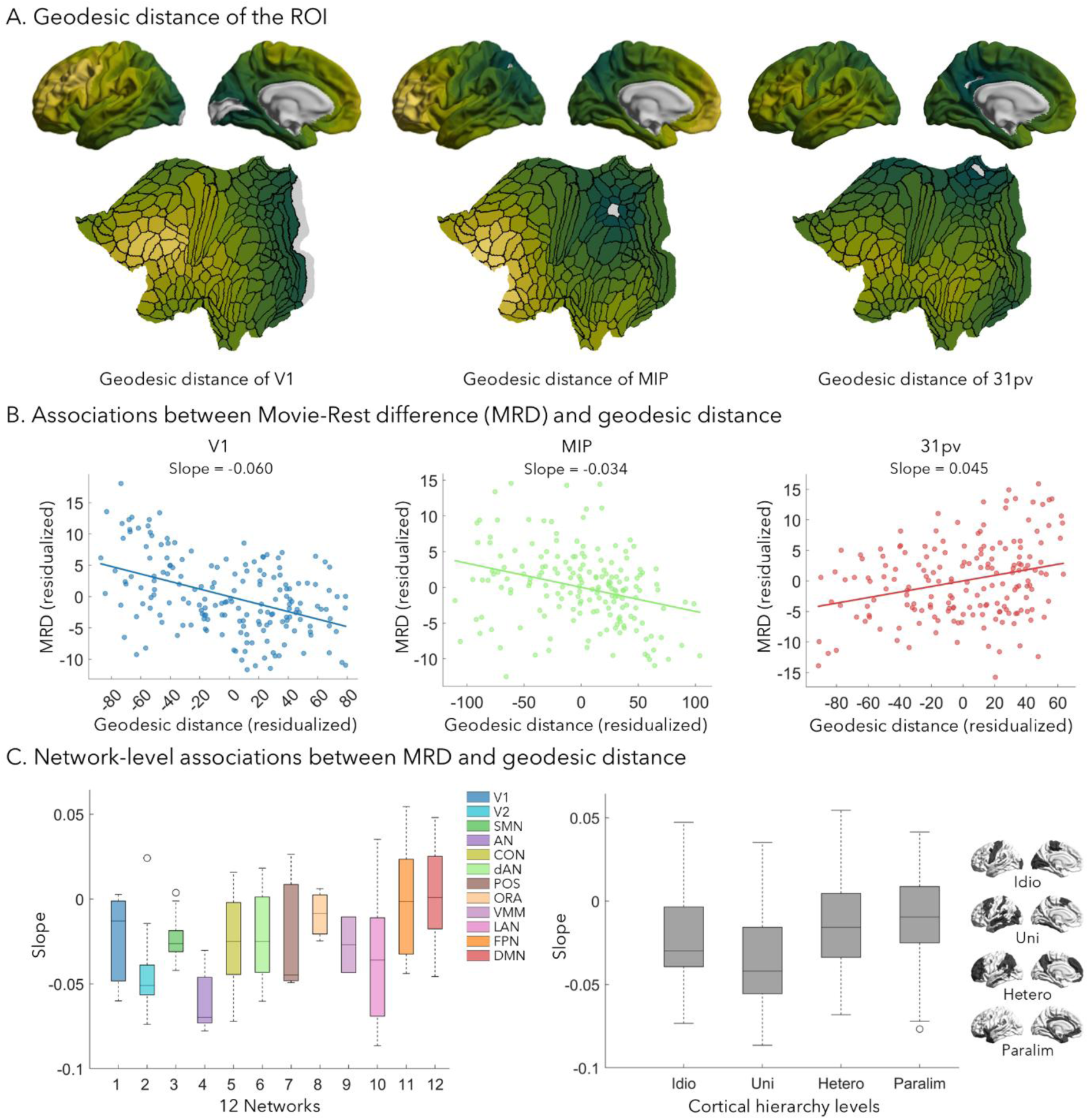
Associations between Movie–Rest functional connectivity differences and cortical geometry. **(A)** Geodesic distance of the ROI. Surface-based geodesic distance maps are shown for three representative regions: V1 (visual network), MIP (dorsal attention network), and 31pv (default mode network). Geodesic distance reflects the shortest path along the cortical surface from each ROI to all other parcels. Warmer colors (e.g. yellow) indicate greater surface-based distance. **(B)** Associations between MRD and geodesic distance. For each ROI, MRD values were regressed against geodesic distance to assess the spatial profile of functional reorganization. V1 and MIP showed negative slopes (–0.060 and –0.034, respectively), indicating stronger connectivity changes in nearby regions during movie-watching. In contrast, 31pv exhibited a positive slope (0.045), suggesting more pronounced reorganization in distant areas. **(C)** Network-level associations between MRD and geodesic distance. Average MRD–GD slopes were computed across all regions within each of the twelve large-scale functional networks. The strongest positive slopes were found in the default mode network (DMN) and frontoparietal network (FPN), indicating long-range reconfiguration. Sensory networks such as the auditory (AN) and visual (VN) networks showed the lowest or negative slopes, reflecting localized changes constrained by cortical proximity.

To investigate the broader topological organization of these state-related changes, we computed the average MRD with all other regions within each of the twelve functional networks (Ji et al., 2019). In addition, to capture hierarchical patterns of reorganization, we grouped cortical parcels into four hierarchical levels idiotypic, unimodal, heteromodal, and paralimbic, based on a taxonomy of cortical processing levels formulated for the primate brain (Mesulam, 1998). We then calculated the mean MRD between each ROI and regions within each of these hierarchy levels. This allowed us to systematically examine how stimulus-driven connectivity modulation is structured along the cortical processing hierarchy, from primary sensory to transmodal association regions.

### Examining the relationship between MRD and cortical geometry

To investigate how MRD in functional connectivity relate to the spatial layout of the cortex, we examined associations with geodesic distance (Ecker et al., 2013; Lariviere et al., 2020; Margulies, Falkiewicz, et al., 2016; Paquola et al., 2020), a measure of anatomical distance along the cortical surface (**Figure 3A**). For each ROI, we computed the geodesic distance between the ROI and all other cortical regions. We then assessed the relationship between group-averaged MRD and geodesic distance across these region pairs using linear regression. To account for potential confounding by overall connectivity strength, we included the mean functional connectivity across rest and movie conditions as a covariate in each regression model. Consequently, we used residualized values of MRD and geodesic distance in subsequent analyses. These residuals may contain negative values. The slope of the MRD–geodesic distance relationship was extracted for each ROI. A positive slope indicates that MRD tends to increase with cortical distance, suggesting greater state-related connectivity changes with remote regions. To explore this spatial relationship across levels of cortical organization, we averaged across the twelve canonical functional networks (Ji et al., 2019) and across the four cortical hierarchy levels (Mesulam, 1998): idiotypic, unimodal, heteromodal, and paralimbic cortex. This analysis allowed us to quantify how the brain’s spatial embedding influences functional reconfiguration in response to naturalistic stimulation.

### Data availability

The MRI data used for the 7T discovery analysis are publicly available through the HCP 1200 Subjects Data Release via the Connectome Coordination Facility (CCF) at https://db.humanconnectome.org (Van Essen et al., 2013). A part of the independent 7T validation dataset is accessible through the Open Science Framework (OSF) at https://osf.io/mhq3f/ (Cabalo et al., 2025).

### Code availability

All preprocessing scripts for structural and functional MRI data are available at https://github.com/MICA-MNI/micapipe (Cruces et al., 2022). Code used for computing functional gradients and performing the main analyses in this study is openly accessible at https://github.com/MICA-MNI/Wang_MovieWatching.

## Results

### Functional connectivity patterns across cortical hierarchies

We analyzed high-field 7T MRI data from 93 unrelated healthy adults (mean age: 29.4 ± 3.3 years; 56 females) from the HCP, and conducted a separate replication analysis using the PNI dataset (see *below*). To characterize whole-brain functional connectivity, we computed Pearson correlation coefficients between regional time series for eight fMRI runs: four during resting-state and four during naturalistic movie-watching. The four rs-fMRI connectivity matrices were averaged to produce a composite Rest connectivity matrix, and the four movie functional connectivity matrices were similarly averaged to yield a Movie connectivity matrix.

To visualize how connectivity patterns vary across the cortical hierarchy, we selected three representative regions of interest (ROIs): V1 in the primary visual cortex (V1), MIP in the dorsal attention network (dAN), and 31pv in the default mode network (DMN) (**Figure 1B**). Spearman correlation analysis revealed high similarity in FC between rest and movie states for the ROIs V1 (rho = 0.77, pspin < 0.001), MIP (rho = 0.89, pspin < 0.001), and 31pv (rho = 0.86, pspin < 0.001). These findings suggest that the overall functional architecture is largely preserved across rest and movie-watching conditions. Functional connectivity patterns for ROIs in the remaining nine networks are presented in **Supplementary Figure 1B**.

### Quantifying functional connectivity differences between movie-watching and rest

To systematically assess functional reorganization across brain states, we computed the Movie– Rest Difference (MRD) in functional connectivity for each ROI using a linear mixed-effects model. This approach accounted for inter-subject variability by modeling participant as a random effect, while brain state (rest *vs*. movie) was treated as a fixed effect (**Figure 2A**). The resulting MRD maps for our three representative ROIs (*i.e.,* V1, MIP, and 31pv) revealed distinct spatial profiles of connectivity reconfiguration (**Figure 2B**). In V1, MRD values were highest in neighboring regions, indicating a localized enhancement of connectivity during movie-watching relative to rest. MIP exhibited modest MRD with no clear spatial preference, suggesting limited task-related reconfiguration. In contrast, 31pv showed a distinct shift in connectivity patterns: MRD was reduced in other DMN regions but increased in distant, non-DMN areas. This pattern implies that movie-watching is associated with reduced intra-DMN coupling while simultaneously enhancing inter-network interactions with externally oriented systems. MRD maps for the remaining nine ROIs are provided in **Supplementary Figure 2**.

To further examine reconfiguration at the network level, we examined average MRD values within twelve predefined large-scale brain networks. For V1, the strongest increases in MRD were observed in the ventral multimodal network (VMN; t = 9.8, p < 0.001), while significant decreases were seen in the somatomotor network (SMN; t = –7.5, p < 0.001) and the cingulo-opercular network (CON; t = –9.7, p < 0.001). MIP similarly showed decreased MRD in the CON (t = –7.7, p < 0.001). Notably, 31pv exhibited increased MRD in the CON (t = 7.3, p < 0.001) and reduced MRD within its own network, the DMN (t = –7.3, p < 0.001), consistent with a redistribution of functional integration away from intrinsic default-mode coupling. When aggregating MRD by hierarchical level based on the Mesulam classification, V1 exhibited the lowest overall MRD, reflecting its local and stable functional role. In contrast, 31pv, a high-level transmodal region, showed elevated MRD values, indicative of greater state-dependent reorganization at the apex of the cortical hierarchy.

### Associations Between MRD and Cortical Geometry

To gain deeper insight into the spatial organization of functional reconfiguration, we investigated how MRDs relate to anatomical distance between regions. We estimated geodesic distance (GD), a surface-based measure of the shortest path between cortical vertices for each ROI (**Figure 3A**). We then examined the association between MRD and GD for each ROI. In V1 and MIP, MRD exhibited negative correlations with GD (V1 slope = –0.060; MIP slope = –0.034), suggesting that movie-watching preferentially increased functional connectivity with nearby regions while reducing long-range interactions. Conversely, 31pv exhibited a positive correlation (slope = 0.045), indicating stronger functional connectivity reorganization with distant regions than with proximal ones (**Figure 3B**). To generalize these observations, we computed MRD–GD slopes for ROIs across the remaining nine functional networks (**Supplementary Figure 3**). Results were aggregated by network and hierarchical classification. Among networks, highest average slopes were observed in DMN and frontoparietal network, both associated with transmodal and integrative processing. In contrast, the auditory network (AN) and VN showed the lowest slopes, reflecting reorganization predominantly constrained to nearby regions (**Figure 3C**). When examining these associations by cortical hierarchy (based on Mesulam, 1998), we found that paralimbic and heteromodal regions exhibited the strongest positive MRD–GD relationships (**Figure 3C**). These findings point to a spatially differentiated mode of reconfiguration, wherein sensory areas become more locally integrated and transmodal regions more globally engaged during naturalistic stimulation.

### Replication in an independent cohort

To assess the reproducibility of our findings, we conducted a replication analysis using 7T MRI data from 30 healthy young adults (mean age = 27.82±3.74 years, 16 females) (Cabalo et al., 2025). Fifteen participants underwent three resting-state and four movie-watching sessions, while the other fifteen completed only one resting-state session. Overall, we observed consistent MRDs in regions MIP and 31pv. While the MRD pattern in V1 exhibited some divergence (**Supplementary Figure 4**), this variability may reflect the complex and region-specific nature of functional reorganization during naturalistic stimulation. Importantly, the associations between MRD and geodesic distance were replicated at both the ROI and network levels, supporting the robustness and reproducibility of the original findings.

## Discussion

Understanding how the brain dynamically reconfigures its functional architecture in response to naturalistic stimulation is a central challenge in cognitive neuroscience. Using high-field 7T fMRI data from two independent datasets, we investigated how functional connectivity reorganizes from rest to movie-watching and how these differences relate to cortical hierarchy and spatial proximity. Our results suggested that while the overall functional architecture is generally preserved across states, there is nevertheless systematic, spatially structured, and hierarchy-dependent reorganization during movie-watching. Specifically, unimodal sensory areas, such as V1, exhibited increased local connectivity and decreased long-range connectivity, whereas transmodal regions, including the DMN, showed the opposite pattern—disrupted local coupling but strengthened distal interactions. This spatially differentiated reconfiguration reveals how the brain adapts its large-scale functional organization to support naturalistic cognition and emphasizes the importance of hierarchical and geometric constraints in shaping state-dependent neural dynamics.

Together with between-network integration, functional specialization is a hallmark of the human cortex (Bassett & Bullmore, 2006; Bullmore & Sporns, 2009; Wang et al., 2025; Wang et al., 2023), and it is known that sensory and transmodal regions differ markedly in cytoarchitecture, connectivity, and temporal integration properties (Margulies, Ghosh, et al., 2016; Mesulam, 1998; Wang et al., 2025). Our observation that V1 becomes more locally integrated during movie-watching aligns with previous reports showing that primary sensory cortices are dominated by short-range connections and support fine-grained, stimulus-specific processing (Felleman & Van Essen, 1991). The increased coupling with nearby visual regions during naturalistic viewing likely reflects an enhancement of local feedforward and lateral interactions driven by bottom-up sensory input. In contrast, transmodal regions such as area 31pv, located in the posterior DMN, showed a different pattern. During movie-watching, these regions exhibited diminished local connectivity alongside increased coupling with distant areas, suggesting a shift from intrinsic, locally organized activity to a more distributed, integrative mode of processing. This pattern may reflect the recruitment of transmodal regions into broader task-related networks during complex, multimodal stimulation (Krienen et al., 2014). By relating functional connectivity changes to cortical GD, we found that the spatial scale of reorganization varied systematically with cortical hierarchy. This gradient suggests that different cortical areas operate at different spatial integration scales depending on their hierarchical level, a view that complements prior evidence of functional gradients across the cortex (Huntenburg et al., 2018; Margulies, Ghosh, et al., 2016).

These spatial patterns are consistent with structural and functional models of cortical organization. Anatomical studies have shown that association cortices are positioned at the apex of cortical hierarchies, with more distributed input–output profiles and greater capacity for multimodal integration (Felleman & Van Essen, 1991; Mesulam, 1998). Functionally, these regions are known to participate in flexible, context-dependent interactions, integrating information over longer timescales and across domains (Hasson et al., 2008). Our findings align with these characterizations and extend them by demonstrating that such flexibility is reflected in shifts in spatial connectivity profiles during naturalistic states. The DMN, for instance, is often described as a task-negative network, but recent evidence suggests that it can support internally oriented cognition and event comprehension across a variety of cognitive contexts (Paquola et al., 2025; Simony et al., 2016; Smallwood et al., 2021; Smallwood et al., 2013; Turnbull et al., 2019). Our observation of reduced intra-DMN coupling and increased long-range connectivity with external systems supports the view that DMN regions adaptively reconfigure to meet task demands, balancing intrinsic and extrinsic processing (Paquola et al., 2025). At the network level, MRD showed similar patterns across canonical systems. Notably, the DMN showed both decreased within-network connectivity and increased connectivity with other networks, underscoring its dual role in introspective and externally engaged cognition (Cole et al., 2014). These network-level differences highlight how functional reorganization is shaped not only by anatomical proximity but also by the functional roles and cognitive demands associated with each system.

The association between MRD and GD across networks further illustrates how spatial embedding constrains functional reorganization (Pang et al., 2023; Pang et al., 2025; Paquola et al., 2020). Transmodal systems, such as the DMN and frontoparietal networks, exhibited steeper positive MRD–GD slopes, indicating increased long-range coordination during movie-watching. In contrast, unimodal networks, such as auditory and visual systems, exhibited flat or negative slopes, consistent with their role in spatially localized processing. This divergence suggests that cognitive state modulates not just the strength of connectivity but also the spatial scale at which regions integrate information. Geodesic distance thus emerges as a key organizing principle, revealing how the physical layout of the cortex interacts with functional demands to shape dynamic network reconfiguration (Margulies, Ghosh, et al., 2016; Oligschläger et al., 2017; Pang et al., 2023; Paquola et al., 2020; Sepulcre et al., 2010; Vázquez-Rodríguez et al., 2019). Methodologically, the use of GD rather than Euclidean distance may allow for a more biologically meaningful quantification of spatial proximity, accounting for the cortical sheet’s folded geometry (Ecker et al., 2013; Lariviere et al., 2020; Paquola et al., 2020; Wang et al., 2023; Weber et al., 2024). This choice was critical for capturing region-specific variations in connectivity modulation, particularly in association cortices where convolutions can obscure true spatial relationships. Moreover, our focus on vertex-wise functional connectivity allowed us to detect fine-grained patterns of reorganization that may be masked in parcellation-based analyses, particularly in regions with high functional heterogeneity (Glasser et al., 2016).

Several limitations should be considered. Although 7T fMRI offers high spatial resolution and signal compared to conventional neuroimaging, it is also more susceptible to artifacts, particularly in inferior frontal and temporal regions. We employed rigorous preprocessing and quality control procedures, but residual noise may still affect some estimates. Additionally, while we interpret MRD as reflecting functional reorganization due to cognitive engagement, non-specific factors such as arousal, attention, or eye movements may also contribute to state differences. Future work incorporating physiological or behavioral monitoring could help disentangle these effects. Furthermore, while our models accounted for individual variation using mixed-effects approaches, the generalizability of findings across subjects and stimuli remains an open question. Individual differences in movie comprehension, engagement, or strategy may modulate reorganization patterns, and studies using different stimuli or tasks could provide complementary insights.

Our findings have broad implications for understanding the flexible architecture of the brain. They support a model in which cortical regions dynamically adjust their integration profiles depending on both hierarchical position and cognitive state. Sensory regions appear to operate under local integration principles during naturalistic stimulation, while transmodal regions engage in distributed communication, potentially supporting event segmentation, mentalizing, or narrative integration (Hasson et al., 2008; Simony et al., 2016). These results extend the concept of functional gradients and suggest that dynamic reconfiguration along these gradients is a core feature of naturalistic cognition. More broadly, the findings emphasize the importance of considering spatial geometry and hierarchical structure when studying functional dynamics, particularly in rich, ecologically valid conditions. In conclusion, this study demonstrates that the spatial organization of functional reconfiguration from rest to movie-watching follows a structured, hierarchy-dependent gradient. Connectivity becomes more localized in sensory cortices and more distributed in association areas, with geodesic distance providing a principled metric for understanding these transformations. By integrating geometric, functional, and network-level perspectives, our results shed light on the principles governing brain-wide communication during naturalistic cognition and highlight the value of high-resolution imaging for revealing the nuanced topography of functional brain dynamics.

## Declarations

## Acknowledgements

Yezhou Wang, Dr. Jordan DeKraker, Dr. Alan Evans, and Dr. Boris Bernhardt were supported by the Helmholtz International BigBrain Analytics and Learning Laboratory (HiBALL). Yezhou Wang was funded by the Fonds de recherche du Québec-Nature and technologies (FRQNT). Dr. Jordan DeKraker was funded through a Natural Sciences and Engineering Research Council Post Doctoral Fellowship and a postdoc fellowship from the Montreal Neurological Institute. Dr. Jessica Royer was supported by a fellowship from the Canadian Institute of Health Research (CIHR). Dr. Sofie Valk was funded by the Max Planck Institute. Dr. Boris Bernhardt furthermore acknowledges research support from the National Science and Engineering Research Council of Canada (NSERC Discovery-1304413), CIHR (FDN-154298, PJT-174995, PJT-191853), SickKids Foundation (NI17-039), BrainCanada (Future-Leaders), Healthy Brains and Healthy Lives, the Centre of Excellence in Epilepsy at the Neuro (CEEN), and the Tier-2 Canada Research Chairs (CRC) program.

## Author Contributions

Y.W. conceptualized the project, designed the methodology, conducted the analyses and drafted the manuscript. J.D., assisted with data processing and analysis. R.R.-C., D.G.C., J.R., A.N., M.S., Y.H., and I.R.L. contributed to data acquisition and preprocessing of the imaging datasets. T.V. provided support for the movie-watching fMRI dataset. R.N.S. and S.L.V. offered guidance on the functional imaging analyses. A.C.E. contributed detailed input on the statistical analyses. B.C.B. co-conceptualized the project, supervised its implementation, and revised the manuscript.

## Ethical statement

We confirm that all relevant ethical regulations were adhered to in the design and conduct of this study.

## References

Alexander-Bloch, A. F., Vértes, P. E., Stidd, R., Lalonde, F., Clasen, L., Rapoport, J.,…Gogtay, N. (2013). The Anatomical Distance of Functional Connections Predicts Brain Network Topology in Health and Schizophrenia. Cerebral Cortex, 23(1), 127–138. 10.1093/cercor/bhr388

Bassett, D. S., & Bullmore, E. (2006). Small-World Brain Networks. The Neuroscientist, 12(6), 512–523. 10.1177/1073858406293182

Bernhardt, B. C., Smallwood, J., Keilholz, S., & Margulies, D. S. (2022). Gradients in brain organization. Neuroimage, 251, 118987. 10.1016/j.neuroimage.2022.118987

Bernhardt, B. C., Valk, S. L., Hong, S.-J., Soulières, I., & Mottron, L. (2025). Autism-related shifts in the brain&#x2019;s information processing hierarchy. Trends in Cognitive Sciences. 10.1016/j.tics.2025.04.008

Biswal, B., Zerrin Yetkin, F., Haughton, V. M., & Hyde, J. S. (1995). Functional connectivity in the motor cortex of resting human brain using echo-planar mri [10.1002/mrm.1910340409]. Magnetic Resonance in Medicine, 34(4), 537-541. 10.1002/mrm.1910340409

Bullmore, E., & Sporns, O. (2009). Complex brain networks: graph theoretical analysis of structural and functional systems. Nature Reviews: Neuroscience, 10(3), 186–198. 10.1038/nrn2575

Cabalo, D. G., Leppert, I. R., Thevakumaran, R., DeKraker, J., Hwang, Y., Royer, J.,…Bernhardt, B. C. (2025). Multimodal precision MRI of the individual human brain at ultra-high fields. Scientific Data, 12(1), 526. 10.1038/s41597-025-04863-7

Cole, Michael W., Bassett, Danielle S., Power, Jonathan D., Braver, Todd S., & Petersen, Steven E. (2014). Intrinsic and Task-Evoked Network Architectures of the Human Brain. Neuron, 83(1), 238–251. 10.1016/j.neuron.2014.05.014

Cole, M. W., Ito, T., Bassett, D. S., & Schultz, D. H. (2016). Activity flow over resting-state networks shapes cognitive task activations. Nature Neuroscience, 19(12), 1718–1726. 10.1038/nn.4406

Coupe, P., Yger, P., Prima, S., Hellier, P., Kervrann, C., & Barillot, C. (2008). An Optimized Blockwise Nonlocal Means Denoising Filter for 3-D Magnetic Resonance Images. IEEE Transactions on Medical Imaging, 27(4), 425–441. 10.1109/TMI.2007.906087

Cox, R. W. (1996). AFNI: Software for Analysis and Visualization of Functional Magnetic Resonance Neuroimages. Computers and Biomedical Research, 29(3), 162–173. 10.1006/cbmr.1996.0014

Cruces, R. R., Royer, J., Herholz, P., Larivière, S., Vos de Wael, R., Paquola, C.,…Bernhardt, B. C. (2022). Micapipe: A pipeline for multimodal neuroimaging and connectome analysis. Neuroimage, 263, 119612. 10.1016/j.neuroimage.2022.119612

Demirtaş, M., Burt, J. B., Helmer, M., Ji, J. L., Adkinson, B. D., Glasser, M. F.,…Murray, J. D. (2019). Hierarchical Heterogeneity across Human Cortex Shapes Large-Scale Neural Dynamics. Neuron, 101(6), 1181–1194.e1113. 10.1016/j.neuron.2019.01.017

DuPre, E., Salo, T., Ahmed, Z., Bandettini, P. A., Bottenhorn, K. L., Caballero-Gaudes, C.,…Handwerker, D. A. (2021). TE-dependent analysis of multi-echo fMRI with tedana. Journal of Open Source Software, 6(66), 3669. 10.21105/joss.03669

Ecker, C., Ronan, L., Feng, Y., Daly, E., Murphy, C., Ginestet, C. E.,…Williams, S. C. (2013). Intrinsic gray-matter connectivity of the brain in adults with autism spectrum disorder. Proceedings of the National Academy of Sciences, 110(32), 13222–13227. 10.1073/pnas.1221880110

Felleman, D. J., & Van Essen, D. C. (1991). Distributed Hierarchical Processing in the Primate Cerebral Cortex. Cerebral Cortex, 1(1), 1–47. 10.1093/cercor/1.1.1-a

Finn, E. S., & Bandettini, P. A. (2021). Movie-watching outperforms rest for functional connectivity-based prediction of behavior. Neuroimage, 235, 117963. 10.1016/j.neuroimage.2021.117963

Finn, E. S., Shen, X., Scheinost, D., Rosenberg, M. D., Huang, J., Chun, M. M.,…Constable, R. T. (2015). Functional connectome fingerprinting: identifying individuals using patterns of brain connectivity. Nature Neuroscience, 18(11), 1664–1671. 10.1038/nn.4135

Fischl, B. (2012). FreeSurfer. Neuroimage, 62(2), 774–781. 10.1016/j.neuroimage.2012.01.021

Fox, M. D., Snyder, A. Z., Vincent, J. L., Corbetta, M., Van Essen, D. C., & Raichle, M. E. (2005). The human brain is intrinsically organized into dynamic, anticorrelated functional networks. Proceedings of the National Academy of Sciences, 102(27), 9673–9678. 10.1073/pnas.0504136102

Glasser, M. F., Coalson, T. S., Robinson, E. C., Hacker, C. D., Harwell, J., Yacoub, E.,…Van Essen, D. C. (2016). A multi-modal parcellation of human cerebral cortex. Nature, 536(7615), 171–178. 10.1038/nature18933

Glasser, M. F., Sotiropoulos, S. N., Wilson, J. A., Coalson, T. S., Fischl, B., Andersson, J. L.,…Jenkinson, M. (2013). The minimal preprocessing pipelines for the Human Connectome Project. Neuroimage, 80, 105–124. 10.1016/j.neuroimage.2013.04.127

Goulas, A., Uylings, H. B. M., & Hilgetag, C. C. (2017). Principles of ipsilateral and contralateral cortico-cortical connectivity in the mouse. Brain Structure and Function, 222(3), 1281–1295. 10.1007/s00429-016-1277-y

Greve, D. N., & Fischl, B. (2009). Accurate and robust brain image alignment using boundary-based registration. Neuroimage, 48(1), 63–72. 10.1016/j.neuroimage.2009.06.060

Haast, R. A. M., Ivanov, D., Formisano, E., & Uludaǧ, K. (2016). Reproducibility and Reliability of Quantitative and Weighted T(1) and T(2)(∗) Mapping for Myelin-Based Cortical Parcellation at 7 Tesla. Frontiers in Neuroanatomy, 10, 112–112. 10.3389/fnana.2016.00112

Hasson, U., Nir, Y., Levy, I., Fuhrmann, G., & Malach, R. (2004). Intersubject Synchronization of Cortical Activity During Natural Vision. Science, 303(5664), 1634–1640. 10.1126/science.1089506

Hasson, U., Yang, E., Vallines, I., Heeger, D. J., & Rubin, N. (2008). A Hierarchy of Temporal Receptive Windows in Human Cortex. the journal of neuroscience, 28(10), 2539. 10.1523/JNEUROSCI.5487-07.2008

Henschel, L., Conjeti, S., Estrada, S., Diers, K., Fischl, B., & Reuter, M. (2020). FastSurfer - A fast and accurate deep learning based neuroimaging pipeline. Neuroimage, 219, 117012. 10.1016/j.neuroimage.2020.117012

Huntenburg, J. M., Bazin, P.-L., & Margulies, D. S. (2018). Large-Scale Gradients in Human Cortical Organization. Trends in Cognitive Sciences, 22(1), 21–31. 10.1016/j.tics.2017.11.002

Jenkinson, M., Beckmann, C. F., Behrens, T. E. J., Woolrich, M. W., & Smith, S. M. (2012). FSL. Neuroimage, 62(2), 782–790. 10.1016/j.neuroimage.2011.09.015

Ji, J. L., Spronk, M., Kulkarni, K., Repovš, G., Anticevic, A., & Cole, M. W. (2019). Mapping the human brain’s cortical-subcortical functional network organization. Neuroimage, 185, 35–57. 10.1016/j.neuroimage.2018.10.006

Krienen, F. M., Yeo, B. T. T., & Buckner, R. L. (2014). Reconfigurable task-dependent functional coupling modes cluster around a core functional architecture. Philosophical Transactions of the Royal Society B: Biological Sciences, 369(1653), 20130526. 10.1098/rstb.2013.0526

Lariviere, S., Weng, Y., Vos de Wael, R., Royer, J., Frauscher, B., Wang, Z.,…Bernhardt, B. C. (2020). Functional connectome contractions in temporal lobe epilepsy: Microstructural underpinnings and predictors of surgical outcome. Epilepsia, 61(6), 1221–1233. 10.1111/epi.16540

Lemieux, L., Salek-Haddadi, A., Lund, T. E., Laufs, H., & Carmichael, D. (2007). Modelling large motion events in fMRI studies of patients with epilepsy. Magnetic Resonance Imaging, 25(6), 894–901. 10.1016/j.mri.2007.03.009

Margulies, D. S., Falkiewicz, M., & Huntenburg, J. M. (2016). A cortical surface-based geodesic distance package for Python. GigaScience, 5(suppl_1). 10.1186/s13742-016-0147-0-q

Margulies, D. S., Ghosh, S. S., Goulas, A., Falkiewicz, M., Huntenburg, J. M., Langs, G.,…Smallwood, J. (2016). Situating the default-mode network along a principal gradient of macroscale cortical organization. Proceedings of the National Academy of Sciences of the United States of America, 113(44), 12574–12579. 10.1073/pnas.1608282113

Marques, J. P., Kober, T., Krueger, G., van der Zwaag, W., Van de Moortele, P.-F., & Gruetter, R. (2010). MP2RAGE, a self bias-field corrected sequence for improved segmentation and T1-mapping at high field. Neuroimage, 49(2), 1271–1281. 10.1016/j.neuroimage.2009.10.002

Mesulam, M. M. (1998). From sensation to cognition. Brain, 121(6), 1013–1052. 10.1093/brain/121.6.1013

Nastase, S. A., Goldstein, A., & Hasson, U. (2020). Keep it real: rethinking the primacy of experimental control in cognitive neuroscience. Neuroimage, 222, 117254. 10.1016/j.neuroimage.2020.117254

Ngo, G. H., Khosla, M., Jamison, K., Kuceyeski, A., & Sabuncu, M. R. (2022). Predicting individual task contrasts from resting-state functional connectivity using a surface-based convolutional network. Neuroimage, 248, 118849. 10.1016/j.neuroimage.2021.118849

Oligschläger, S., Huntenburg, J. M., Golchert, J., Lauckner, M. E., Bonnen, T., & Margulies, D. S. (2017). Gradients of connectivity distance are anchored in primary cortex. Brain Structure and Function, 222(5), 2173–2182. 10.1007/s00429-016-1333-7

Pang, J. C., Aquino, K. M., Oldehinkel, M., Robinson, P. A., Fulcher, B. D., Breakspear, M., & Fornito, A. (2023). Geometric constraints on human brain function. Nature, 618(7965), 566–574. 10.1038/s41586-023-06098-1

Pang, J. C., Robinson, P. A., Aquino, K. M., Levi, P. T., Holmes, A., Markicevic, M.,…Fornito, A. (2025). Geometric influences on the regional organization of the mammalian brain. bioRxiv, 2025.2001.2030.635820. 10.1101/2025.01.30.635820

Paquola, C., Garber, M., Frässle, S., Royer, J., Zhou, Y., Tavakol, S.,…Bernhardt, B. C. (2025). The architecture of the human default mode network explored through cytoarchitecture, wiring and signal flow. Nature Neuroscience, 28(3), 654–664. 10.1038/s41593-024-01868-0

Paquola, C., Seidlitz, J., Benkarim, O., Royer, J., Klimes, P., Bethlehem, R. A. I.,…Bernhardt, B. C. (2020). A multi-scale cortical wiring space links cellular architecture and functional dynamics in the human brain. PLoS Biology, 18(11), e3000979. 10.1371/journal.pbio.3000979

Paquola, C., Vos De Wael, R., Wagstyl, K., Bethlehem, R. A. I., Hong, S.-J., Seidlitz, J.,…Bernhardt, B. C. (2019). Microstructural and functional gradients are increasingly dissociated in transmodal cortices. PLoS Biology, 17(5), e3000284. 10.1371/journal.pbio.3000284

Power, Jonathan D., Cohen, Alexander L., Nelson, Steven M., Wig, Gagan S., Barnes, Kelly A., Church, Jessica A.,…Petersen, Steven E. (2011). Functional Network Organization of the Human Brain. Neuron, 72(4), 665–678. 10.1016/j.neuron.2011.09.006

Raichle, M. E., MacLeod, A. M., Snyder, A. Z., Powers, W. J., Gusnard, D. A., & Shulman, G. L. (2001). A default mode of brain function. Proceedings of the National Academy of Sciences, 98(2), 676–682. 10.1073/pnas.98.2.676

Salimi-Khorshidi, G., Douaud, G., Beckmann, C. F., Glasser, M. F., Griffanti, L., & Smith, S. M. (2014). Automatic denoising of functional MRI data: Combining independent component analysis and hierarchical fusion of classifiers. Neuroimage, 90, 449–468. 10.1016/j.neuroimage.2013.11.046

Samara, A., Eilbott, J., Margulies, D. S., Xu, T., & Vanderwal, T. (2023). Cortical gradients during naturalistic processing are hierarchical and modality-specific. Neuroimage, 271, 120023. 10.1016/j.neuroimage.2023.120023

Satterthwaite, T. D., Elliott, M. A., Gerraty, R. T., Ruparel, K., Loughead, J., Calkins, M. E.,…Wolf, D. H. (2013). An improved framework for confound regression and filtering for control of motion artifact in the preprocessing of resting-state functional connectivity data. Neuroimage, 64, 240–256. 10.1016/j.neuroimage.2012.08.052

Sepulcre, J., Liu, H., Talukdar, T., Martincorena, I., Yeo, B. T. T., & Buckner, R. L. (2010). The Organization of Local and Distant Functional Connectivity in the Human Brain. PLoS Computational Biology, 6(6), e1000808. 10.1371/journal.pcbi.1000808

Simony, E., Honey, C. J., Chen, J., Lositsky, O., Yeshurun, Y., Wiesel, A., & Hasson, U. (2016). Dynamic reconfiguration of the default mode network during narrative comprehension. Nature Communications, 7(1), 12141. 10.1038/ncomms12141

Smallwood, J., Bernhardt, B. C., Leech, R., Bzdok, D., Jefferies, E., & Margulies, D. S. (2021). The default mode network in cognition: a topographical perspective. Nature Reviews Neuroscience, 22(8), 503–513. 10.1038/s41583-021-00474-4

Smallwood, J., Tipper, C., Brown, K., Baird, B., Engen, H., Michaels, J. R.,…Schooler, J. W. (2013). Escaping the here and now: Evidence for a role of the default mode network in perceptually decoupled thought. Neuroimage, 69, 120–125. 10.1016/j.neuroimage.2012.12.012

Smith, S. M., Fox, P. T., Miller, K. L., Glahn, D. C., Fox, P. M., Mackay, C. E.,…Beckmann, C. F. (2009). Correspondence of the brain’s functional architecture during activation and rest. Proceedings of the National Academy of Sciences, 106(31), 13040. 10.1073/pnas.0905267106

Sonkusare, S., Breakspear, M., & Guo, C. (2019). Naturalistic Stimuli in Neuroscience: Critically Acclaimed. Trends in Cognitive Sciences, 23(8), 699–714. 10.1016/j.tics.2019.05.004

Sydnor, V. J., Larsen, B., Bassett, D. S., Alexander-Bloch, A., Fair, D. A., Liston, C.,…Satterthwaite, T. D. (2021). Neurodevelopment of the association cortices: Patterns, mechanisms, and implications for psychopathology. Neuron, 109(18), 2820–2846. 10.1016/j.neuron.2021.06.016

Tavor, I., Jones, O. P., Mars, R. B., Smith, S. M., Behrens, T. E., & Jbabdi, S. (2016). Task-free MRI predicts individual differences in brain activity during task performance. Science, 352(6282), 216–220. 10.1126/science.aad8127

Turnbull, A., Wang, H. T., Murphy, C., Ho, N. S. P., Wang, X., Sormaz, M.,…Smallwood, J. (2019). Left dorsolateral prefrontal cortex supports context-dependent prioritisation of off-task thought. Nature Communications, 10(1), 3816. 10.1038/s41467-019-11764-y

Tustison, N. J., Avants, B. B., Cook, P. A., Zheng, Y., Egan, A., Yushkevich, P. A., & Gee, J. C. (2010). N4ITK: Improved N3 Bias Correction. IEEE Transactions on Medical Imaging, 29(6), 1310–1320. 10.1109/TMI.2010.2046908

Van Essen, D. C., Smith, S. M., Barch, D. M., Behrens, T. E. J., Yacoub, E., & Ugurbil, K. (2013). The WU-Minn Human Connectome Project: An overview. Neuroimage, 80, 62–79. 10.1016/j.neuroimage.2013.05.041

Vanderwal, T., Eilbott, J., & Castellanos, F. X. (2019). Movies in the magnet: Naturalistic paradigms in developmental functional neuroimaging. Developmental Cognitive Neuroscience, 36, 100600. 10.1016/j.dcn.2018.10.004

Vanderwal, T., Eilbott, J., Finn, E. S., Craddock, R. C., Turnbull, A., & Castellanos, F. X. (2017). Individual differences in functional connectivity during naturalistic viewing conditions. Neuroimage, 157, 521–530. 10.1016/j.neuroimage.2017.06.027

Vázquez-Rodríguez, B., Suárez, L. E., Markello, R. D., Shafiei, G., & Paquola, C. (2019). Gradients of structure–function tethering across neocortex. Proceedings of the National Academy of Sciences, 116(42), 21219–21227.

Wang, Y., Eichert, N., Paquola, C., Rodriguez-Cruces, R., DeKraker, J., Royer, J.,…Bernhardt, B. C. (2025). Multimodal gradients unify local and global cortical organization. Nature Communications, 16(1), 3911. 10.1038/s41467-025-59177-4

Wang, Y., Royer, J., Park, B.-y., Vos de Wael, R., Larivière, S., Tavakol, S.,…Bernhardt, B. C. (2023). Long-range functional connections mirror and link microarchitectural and cognitive hierarchies in the human brain. Cerebral Cortex, 33(5), 1782–1798. 10.1093/cercor/bhac172

Weber, C. F., Kebets, V., Benkarim, O., Lariviere, S., Wang, Y., Ngo, A.,…Bernhardt, B. C. (2024). Contracted functional connectivity profiles in autism. Molecular Autism, 15(1), 38. 10.1186/s13229-024-00616-2

Wei, W., Benn, R. A., Scholz, R., Shevchenko, V., Klatzmann, U., Alberti, F.,…Margulies, D. S. (2024). A function-based mapping of sensory integration along the cortical hierarchy. Communications Biology, 7(1), 1593. 10.1038/s42003-024-07224-z

Yeo, B. T. T., Krienen, F. M., Sepulcre, J., Sabuncu, M. R., Lashkari, D., Hollinshead, M.,…Buckner, R. L. (2011). The organization of the human cerebral cortex estimated by intrinsic functional connectivity. Journal of Neurophysiology, 106(3), 1125–1165. 10.1152/jn.00338.2011

