## Supplemental Information for "State dependent shifts in large scale functional topographies"

* Yezhou Wang

* Boris C. Bernhardt

**Figures**

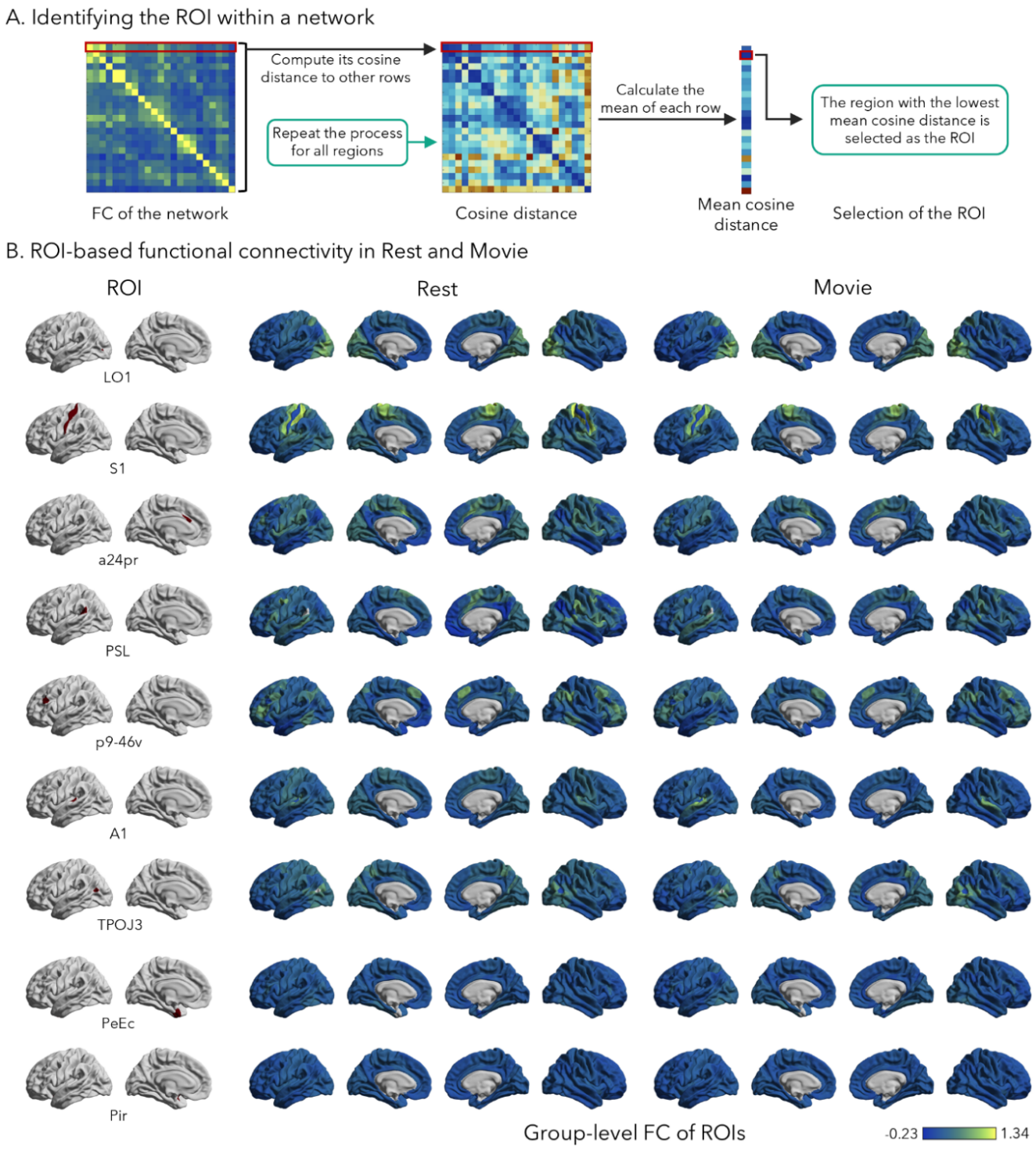

**Supplementary Figure 1.** **(A)** Identification of ROIs within each network. For every region within a given network, we calculated its mean cosine distance to all other regions in the same network. The region with the lowest mean cosine distance was selected as the representative region of interest (ROI), reflecting maximal similarity to other network nodes. **(B)** ROI-based functional connectivity in Rest and Movie. ROI selection was performed for all twelve networks. Results for the V1 (visual network), MIP (dorsal attention network), and 31pv (default mode network) are presented in Figure 1. The group-level functional connectivity (FC) profiles of the selected ROIs during rest and movie-watching conditions for the remaining nine networks are shown in the right panel.

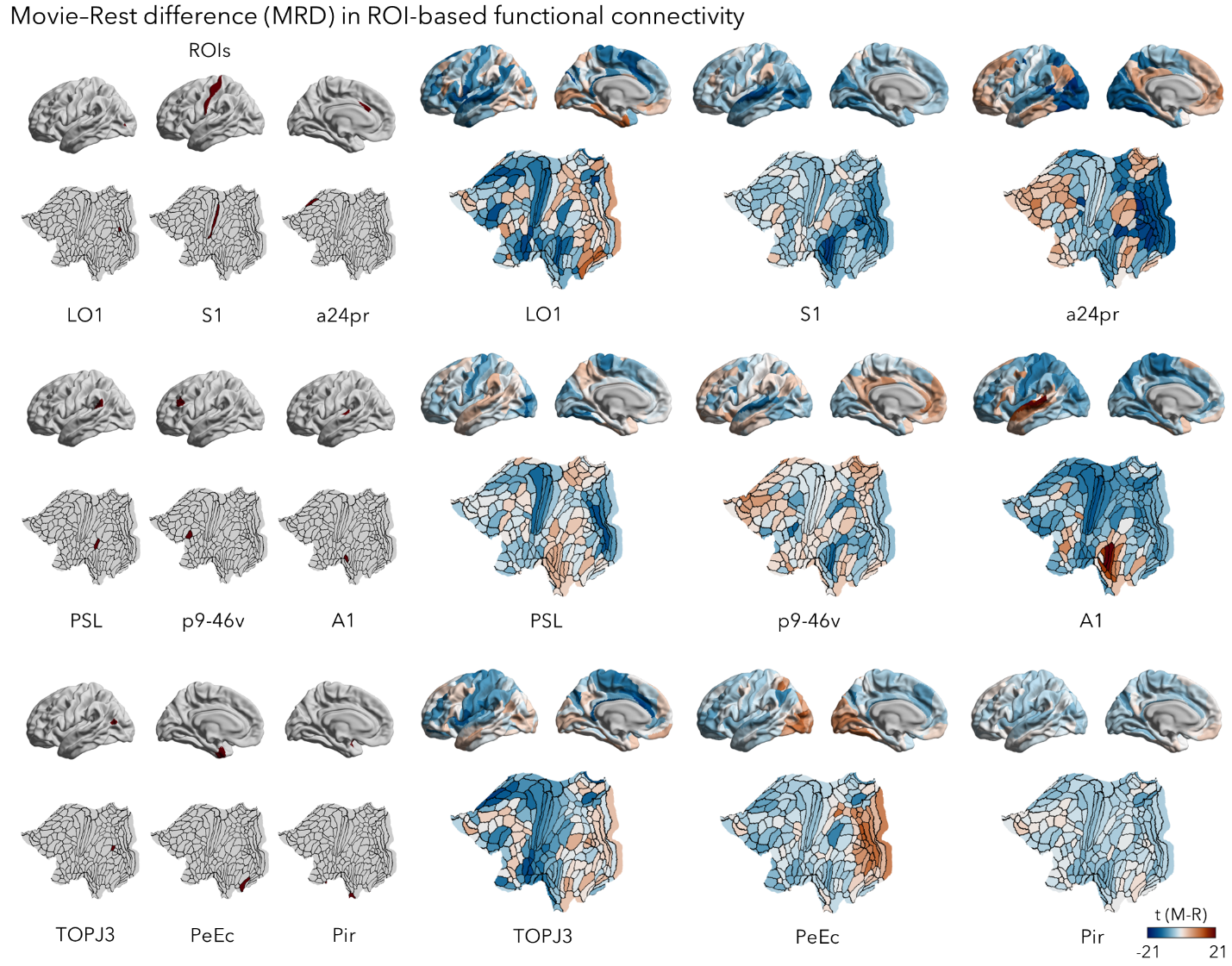

**Supplementary Figure 2.** MRD in ROI-based functional connectivity for the nine remaining networks. The selected ROIs (highlighted in red) for each network are shown in the left panel. The right panel displays the corresponding MRD maps, where warmer colors indicate parcels with stronger functional connectivity to the ROI during movie-watching compared to rest. Results for the V1 (visual network), MIP (dorsal attention network), and 31pv (default mode network) are shown in *Figure 2B.*

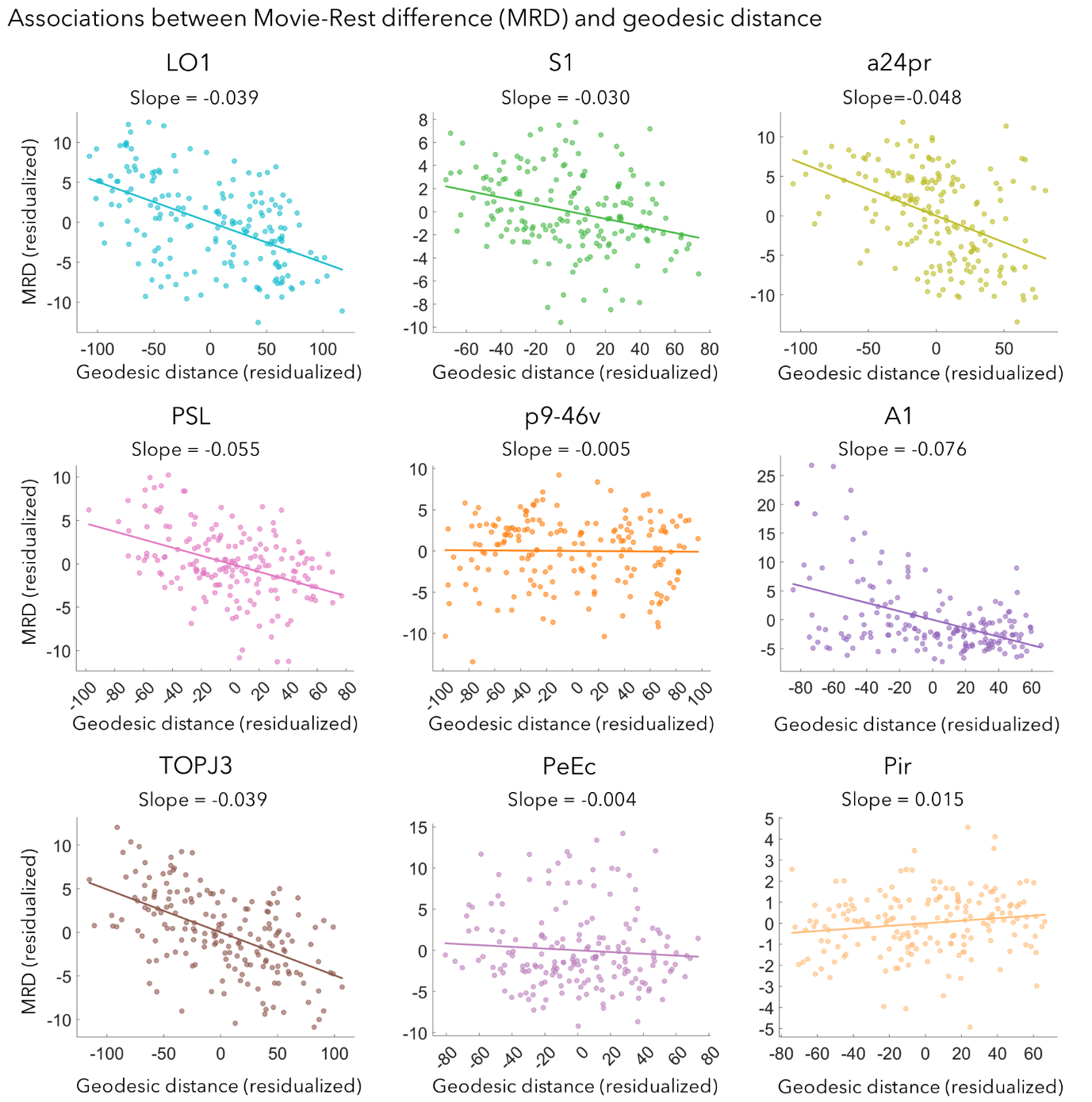

**Supplementary Figure 3.** Associations between MRD and geodesic distance for the nine remaining ROIs. For each ROI, MRD values were regressed against geodesic distance to assess the spatial profile of functional reorganization. Results for V1 (visual network), MIP (dorsal attention network), and 31pv (default mode network) are presented in *Figure 3B*.

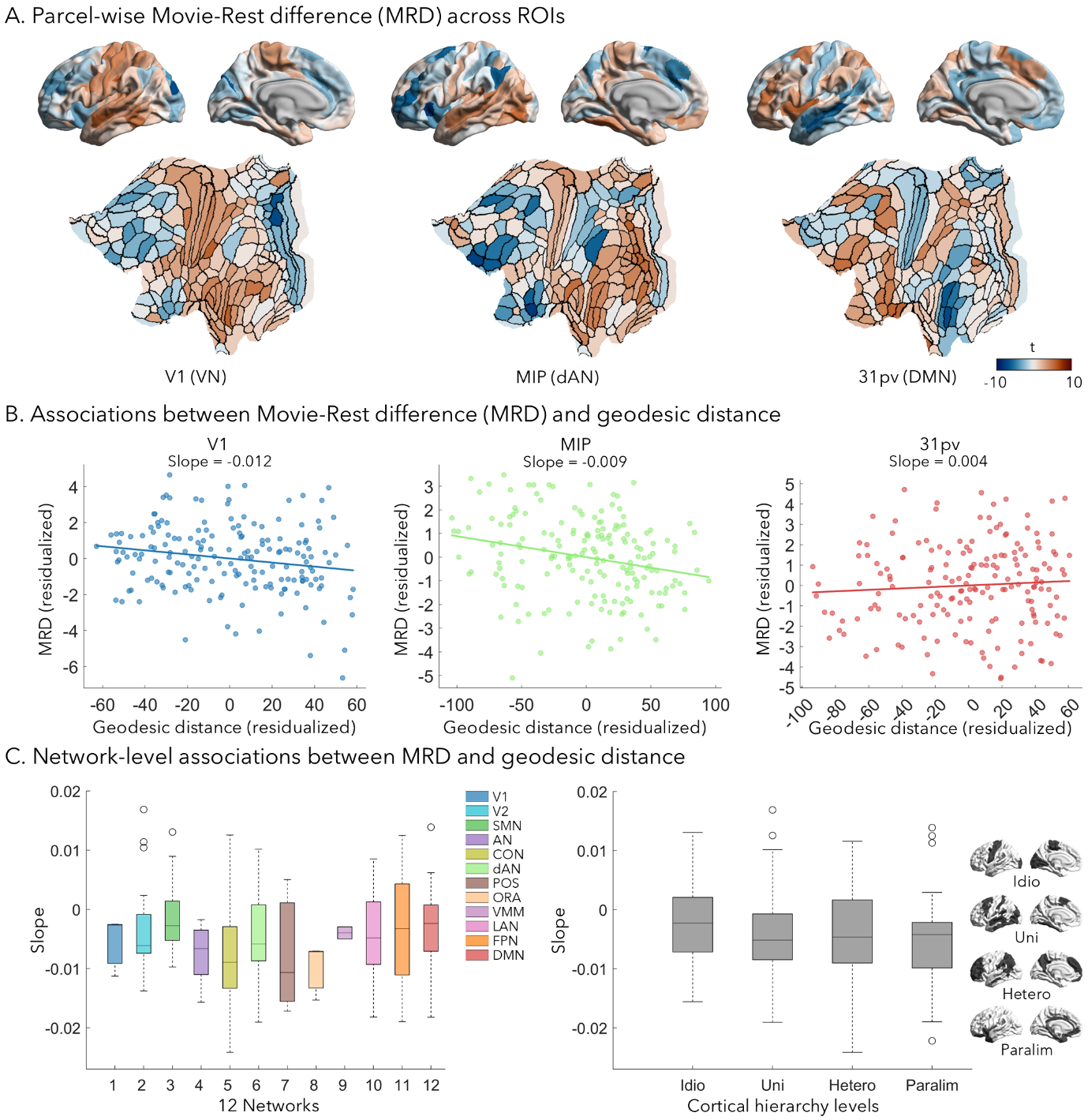

**Supplementary Figure 4. Replication analysis with the PNI 7T dataset.** (A) Parcel-wise MRD across ROIs. Parcel-wise MRD maps are displayed for three representative regions: V1 (visual network, VN), MIP (dorsal attention network, dAN), and 31pv (default mode network, DMN). Warmer colors indicate parcels with stronger functional connectivity to the ROI during movie-watching compared to rest. (B) Associations between MRD and geodesic distance. (C) Network-level associations between MRD and geodesic distance. Average MRD–GD slopes were computed across all regions within each of the twelve large-scale functional networks.
